# Super-Resolution Label-free Volumetric Vibrational Imaging

**DOI:** 10.1101/2021.01.08.425961

**Authors:** Chenxi Qian, Kun Miao, Li-En Lin, Xinhong Chen, Jiajun Du, Lu Wei

**Author notes:** These authors contributed equally: Chenxi Qian, Kun Miao.

## Abstract

Innovations in high-resolution optical imaging have allowed visualization of nanoscale biological structures and connections. However, super-resolution fluorescence techniques, including both optics-oriented and sample-expansion based, are limited in quantification and throughput especially in tissues from photobleaching or quenching of the fluorophores, and low-efficiency or non-uniform delivery of the probes. Here, we report a general sample-expansion vibrational imaging strategy, termed VISTA, for scalable label-free high-resolution interrogations of protein-rich biological structures with resolution down to 82 nm. VISTA achieves decent three-dimensional image quality through optimal retention of endogenous proteins, isotropic sample expansion, and deprivation of scattering lipids. Free from probe-labeling associated issues, VISTA offers unbiased and high-throughput tissue investigations. With correlative VISTA and immunofluorescence, we further validated the imaging specificity of VISTA and trained an image-segmentation model for label-free multi-component and volumetric prediction of nucleus, blood vessels, neuronal cells and dendrites in complex mouse brain tissues. VISTA could hence open new avenues for versatile biomedical studies.

---

Our knowledge of biology is significantly advanced by the development of optical imaging techniques. They reveal multi-dimensional spatial information that is crucial for understanding numerous functions and mechanisms in complex environments. At subcellular levels, super-resolution fluorescence microscopy techniques, including instrument-based stimulated emission depletion (STED) microscopy, photo-activated localization microscopy (PALM) and stochastic optical reconstruction microscopy (STORM)^1,2^, have been devised to overcome the diffraction limit barrier and allow optical visualization of previously unresolvable structures with nanometer resolution. Moreover, sample-oriented strategies, by physically expanding specimens embedded in swellable polymer hydrogels, have made a significant impact on high-resolution imaging of versatile samples^3^. For example, expansion microscopy (ExM) typically achieves a four-fold resolution enhancement using conventional fluorescence microscopes with isotropic sample expansion^4,5^.

Despite their wide applications for uncovering unknown biological events through fine structural and functional characterizations, the super-resolution fluorescence microscopy has a few fundamental limits that originate from the requirement of fluorophore labeling. First, photobleaching and decay of the fluorophores make these techniques less ideal for repetitive and quantitative examinations of target structures^1–5^. This is especially problematic when the sample specimen is limited (e.g., clinical samples). Second, immunofluorescence, the widely used strategy for visualizing various proteins without genetic manipulation, poses serious issues of prolonged sample preparations and inhomogeneous antibody labeling in intact tissues^6^. This is due to the slow diffusion of large antibodies into the tissues and the depletion of probes on the surface^7^. To circumvent these fluorophore-associated challenges, we seek to have a superresolution imaging modality that does not require labels.

Complementary to fluorescence, Raman microscopy targets the specific vibrational motions and maps out the distribution of chemical-bond specific structures and molecules inside live biological systems in a label-free or minimum labeling fashion. In particular, nonlinear stimulated Raman scattering (SRS) has been proven to be a highly successful optical imaging strategy for label-free or tiny-label imaging of biological samples with resolution and speed similar to those of fluorescence^8,9^. For example, SRS imaging targeting the methyl group (i.e. CH_3_) vibrations from endogenous proteins at 2940 cm^-1^ (**Fig. 1a and S1**) has been demonstrated for visualizing proteins-rich structures with submicron resolution at a speed up to video rate in live animals^10^. In principle, implementation of super-resolution Raman imaging could bypass the need of and hence the issues from fluorophore labeling. Despite extensive efforts, this goal has remained challenging. Strategies including excitation saturation, signal suppression with a donut beam or structural illumination have been reported ^11–15^. However, they rely on additional specialized optics and the resolution enhancement is only 1.7 times on biological samples^13–15^. While such optics-based strategies have been heavily explored, there are no known efforts from the perspective of engineering samples for label-free super-resolved Raman imaging.

**Figure 1.**
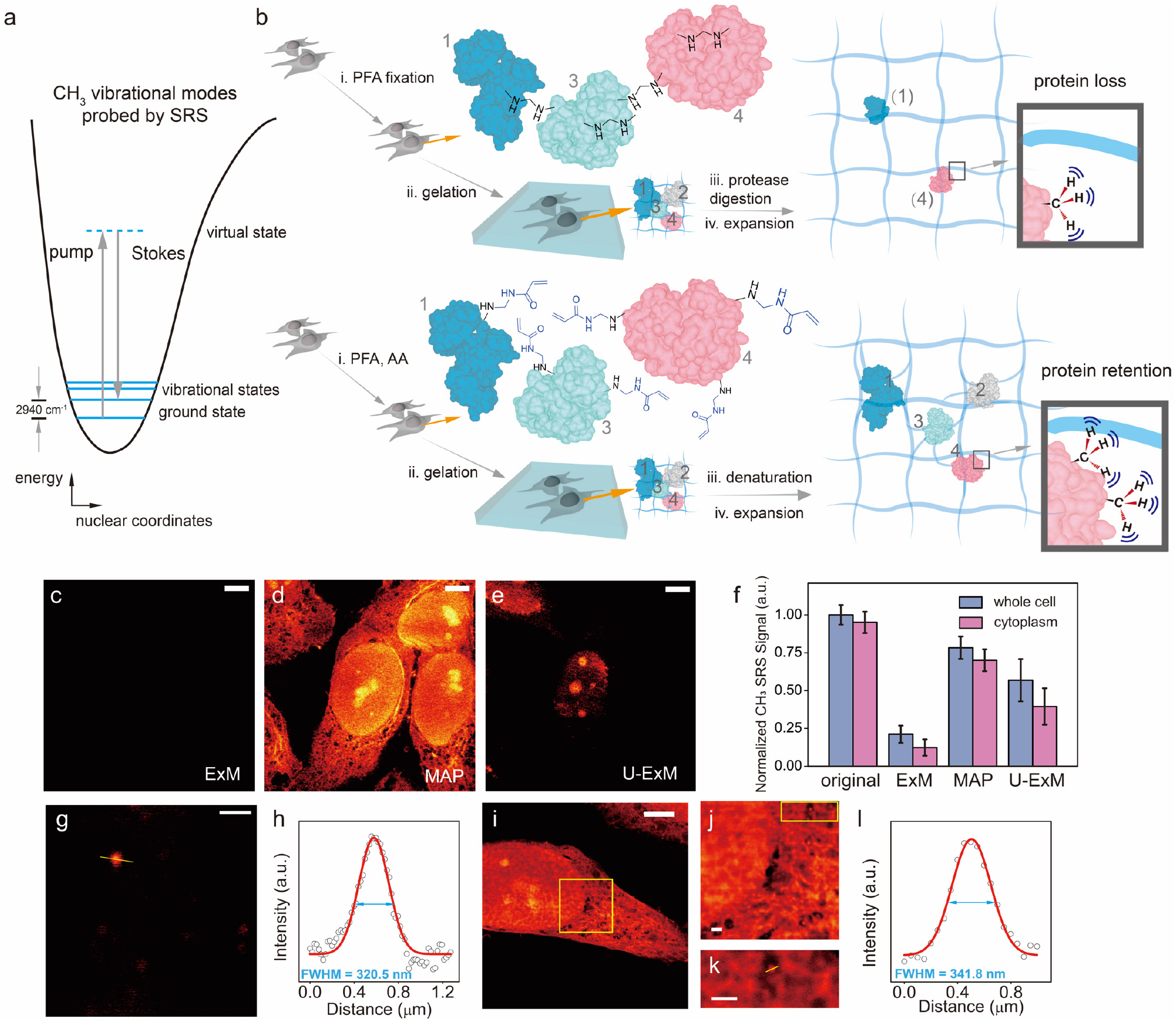
High-resolution label-free vibrational imaging of expanded and protein-retained samples. **a**, Energy scheme for SRS probing of CH_3_ vibrational motion at 2940 cm^-1^. More details in Fig. S1. **b**, Comparison of protein retention (i.e. the methyl groups, CH_3_, from proteins) for SRS imaging between ExM (top) and MAP (bottom) based sample-hydrogel embedding procedures following different fixation, hybridization and homogenization chemistries. **c-e**. SRS imaging of CH_3_ at 2940 cm^-1^ for expanded HeLa cells following ExM, MAP and U-ExM sample treatment under the same intensity scale. Scale bars: 20 μm. **f**, Quantification of proteins retention levels by comparing average CH_3_ signals in expanded cells after ExM, MAP and U-ExM procedures with that from unprocessed HeLa cells (original). CH_3_ signals in expanded cells were scaled back with the average expansion ratios for comparison. Data shown as mean ± SD. **g-h**, Quantification of SRS resolution by imaging the C-H vibration at 3050 cm^-1^ from a representative 100 nm polystyrene bead (g) and fitting its cross-section profile (h). Scale bar: 1 μm. **i-l**, Fitted VISTA imaging cross-section profile (l) from a small structural feature (k) of expanded HeLa cells (i-k, j, k are zoom-in views from the boxed regions in i, j respectively). Scale bars: 30 μm in (i), 2 μm in j-k. For processed samples, the length scale is in terms of distance after expansion.

Here, we report a super-resolution label-free vibrational imaging strategy in cells and tissues that couples the sensitive SRS microscopy with recent sample-treatment innovations. We term this strategy *V*ibrational *I*maging of *S*welled *T*issues and *A*nalysis (VISTA). We embed biological samples in polymer hydrogels, expand the sample-hydrogel hybrid in water, and target the vibrational motion of retained CH_3_ groups from endogenous proteins by SRS for visualization. Our devised strategy possesses a few desirable features. First, compared to fluorescence imaging, VISTA avoids any label-associated issues, allowing uniform imaging and a much higher throughput especially in tissues. Second, compared to optics-based Raman imaging, VISTA is easy to implement without any additional instrument and achieves an unprecedented Raman resolution down to 82 nm on biological samples. Importantly, it allows high-resolution imaging deep into the tissues^16^, a common limit shared by all instrument-based super-resolution microscopy. Third, with further implementation of a convolutional neural network (CNN) for image segmentation^17^, VISTA could offer specific, multi-component, and volumetric imaging in complex tissues with quality similar to that of fluorescence.

As a first step to establish VISTA, we asked whether the sample expansion strategy is compatible with SRS microscopy. We embedded HeLa cells in a polymer gel following the widely-used ExM protocol^4,5^ (**Fig. 1b, top**), which involves paraformaldehyde (PFA) fixation, gelation and sample homogenization through protease digestion. We then performed SRS imaging to visualize hydrogel-retained endogenous proteins in expanded cells at 2940 cm^-1^ (i.e. the CH_3_ channel). However, almost no CH_3_ contrast could be detected (**Fig. 1c, ExM and Fig. S2a**), indicating an extensive loss of proteins or protein fragments under strong protease digestion (i.e., proteinase K)—a known issue in ExM^5^. Indeed, our quantification in CH_3_ channel showed that the protein loss could reach 79% (**Fig. 1f**), consistent with a recent fluorescence analysis^18^. With such a high protein content loss and an approximate 64-fold signal dilution due to 4-fold isotropic sample expansion, SRS signals are therefore diminished. We asked whether reducing digestion time or altering to a milder protease (e.g. Lys-c)^5^ would help retain SRS signals. Unfortunately, proteinase K already significantly digests the protein network even within 30 min (**Fig. S2**). Further reducing the digestion time or changing the protease comes at the expense of low expansion ratios and sample distortion after expansion due to incomplete homogenization of PFA-crosslinked protein networks.

The key to VISTA is to preserve the maximum level of proteins for SRS imaging while achieving optimal homogenization for isotropic sample expansion. Since protease digestion is only required when extensive intra- and inter- protein crosslinking arises from PFA fixation^19^, we resorted to using magnified analysis of proteome (MAP)^20^, an alternative sample-hydrogel hybridization protocol. MAP significantly reduces such PFA-induced protein crosslinking by introducing high concentration of acrylamide (AA) together with PFA fixation so that the excess AA will react with and hence quench the reactive methylols formed by protein-PFA reaction^20^ (**Fig. 1b, bottom**). The subsequent sample homogenization is achieved by protein denaturation instead of protease digestion. With optimizations of incubation time, SRS imaging of CH_3_ showed clear cellular structures with a decent signal-to-noise ratio in expanded cells (**Fig. 1d, MAP**). Characterizations of cell sizes before and after expansion yielded an average expansion ratio of 4 fold, close to that reported through fluorescence^20^. Further quantification of the average CH_3_ signal from MAP- processed cells compared to that from unprocessed cells indeed confirmed that proteins were largely preserved (**Fig. 1f**). Here, the slightly lowered CH_3_ intensity from MAP was likely due to the removal of lipids (**Fig. S3**). Recently, an ultrastructure expansion microscopy (U-ExM) was reported to image ultrastructures with decreased concentration of AA^21^. However, our analysis demonstrated that lowering AA concentration would still lead to a loss of rather significant portion of proteins (**Fig. 1e**, U-ExM and **Fig. 1f**, with an additional loss of 21% in whole cells and 32% in cytoplasm compared to MAP). We therefore concluded that MAP-based sample embedding protocol allows high-quality VISTA imaging of protein-rich subcellular structures that were not clearly identifiable with the normal SRS imaging resolution (**Fig. 1d, MAP vs Fig. S1b,** e.g. the sub-structure of nucleoli and the network in cytosol).

Since the signal of VISTA comes from the CH_3_ channel where the spectral crosstalk of other vibrations might exist, we next examined possible background contributions from both CH2 vibrations of the hydrogel and the O-H stretch of the water. We compared the background sizes and spatial distributions by replacing normal hydrogel monomer and water with their deuterated correspondents respectively. Our data showed that the background introduced from each component can be largely minimized through the deuteration strategy (**Fig. S4**) and the lowering of the signals were uniform throughout the cellular samples, implying that the background is constant and does not introduce any heterogeneous imaging features. After optimizing and confirming the sample processing and imaging conditions, we then aimed to determine the achievable resolution of VISTA. We performed regular SRS imaging on 100 nm polystyrene beads at 3050 cm^-1^ for C-H bonds with a NA=1.05 objective (**Fig. 1g**). We obtained a fitted image cross section with a full width at half maximum (FWHM) of 320 nm, which designates a resolution of 382 nm of the SRS microscope by Rayleigh criterion after deconvolution of the bead object function (**Fig. 1h)**. These data indicate that VISTA offers a 96 nm effective resolution after a 4-fold isotropic sample expansion. We also confirmed a similar level of resolution of VISTA on small structural features within HeLa cells (**Fig. 1i-l**). The effective resolution could be further pushed to an unprecedented 82 nm with a higher-NA objective (i.e. NA=1.2) **(Fig. S5).**

With isotropic sample expansion, VISTA also provides sharp three-dimensional (3D) views of cellular morphology and subcellular structures (**Fig. S6a-b**). In addition to imaging normal cell states, we applied VISTA to visualizing structural changes of HeLa cells throughout mitosis in metaphase, anaphase, telophase and interphase (**Fig. 2a-d**). VISTA images clearly resolved fine structures including cytosolic inner networks, small membrane protrusions of microvilli (**Fig. 2a-d, white arrowed**) and protein-rich contractile ring and midbody (**Fig. 2d, blue arrowed**). In addition, we observed an interesting change of protein contents during mitosis. The relative level of chromosome-associated proteins decreases as the nuclear envelopes disintegrate. This is evidenced by the dark region in cells (**Fig. 2a-c)**, which designates chromosome structures, confirmed by DAPI fluorescence stain (**Fig. S7**). The protein level then increases as the nuclear envelopes reform at the end of the mitosis in telophase (**Fig. 2d, green arrowed**). The 3D views of the cells are shown in **Fig. S6c-d**. These data imply that VISTA is capable of performing quantitative total protein analysis at subcellular compartments.

**Figure 2.**
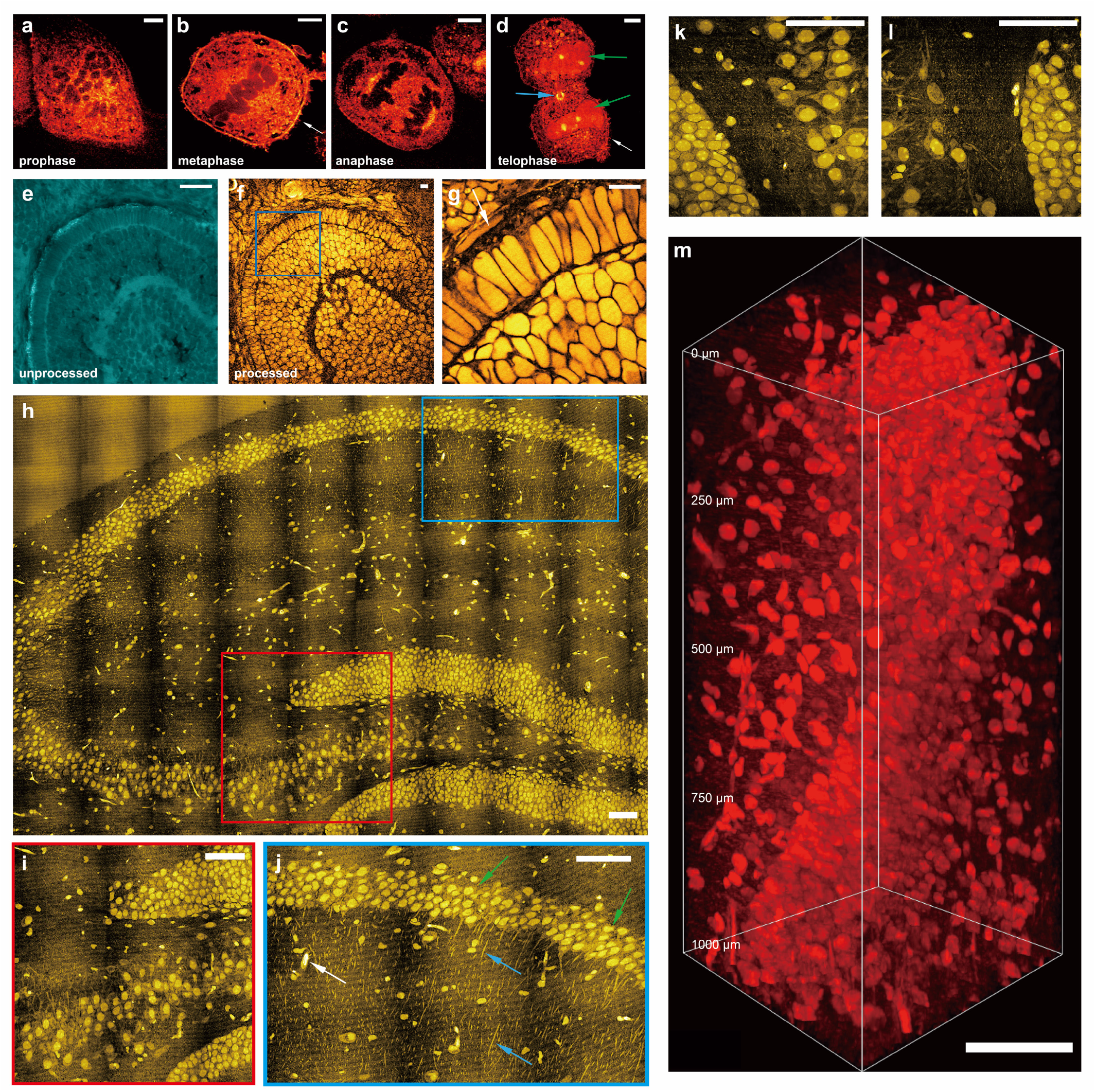
Super-resolution three-dimensional VISTA imaging of cells and tissues. **a-d**, Mitotic HeLa cells in prophase (a), metaphase (b), anaphase (c) and telophase (d). **e-g**, zebrafish embryonic retina before processing and after expansion: unprocessed (e), expanded (f), and zoom-in view from the boxed region in f (g). **h**, A mosaic VISTA image of a hippocampal tissue. **i-l**, zoomed-in high-resolution view from color- boxed areas (i, red box; j, blue box) and selected regions in h. Representative neuronal cell bodies, neuronal processes and likely blood vessel cross-sections are indicated by green, blue and white arrows respectively in j. **m**, 3D volume VISTA imaging of a hippocampal tissue throughout 1000 μm depth. Scale bars: 20 μm (a-g) and 200 μm (h-m). For processed samples, the length scale is in terms of distance after expansion.

Apart from visualizing fine subcellular structures, VISTA is well-suited for tissue imaging. We first demonstrated VISTA on the optical transparent zebrafish embryos, in particular the cone and rod photoreceptors in the outer segment (OS), important model systems for understanding visual perception^22,23^. Comparisons of CH_3_ images before (**Fig. 2e**) and after (**Fig. 2f**) sample embedding and expansion from the same area presented a clear change of contrast due to lipid removal, which allowed us to unambiguously image protein-rich structures, e.g., the retinal pigment epithelium (**Fig. 2g,** arrow indicated). Our data also confirmed a similar level of sample expansion ratio on tissues compared to that of cells (**Fig. S8**). We then aimed to implement VISTA on the much more scattering and complex mouse brain tissues, especially for the hippocampus where characterizations of intricate structural relationship are important for functional understanding of a series of physiological (e.g. memory formation) and pathological (e.g. neurodegenerative diseases) events^24,25^. Our mosaic VISTA image on hippocampus (**Fig. 2h**) reveals clear and specific contrast from neuronal cell bodies, processes and also likely blood vessel cross-sections at various locations (**Fig. 2i-l, green, blue and white arrows indicated, respectively, in Fig. 2j**). All these features are virtually indistinguishable in regular SRS images due to a much lower resolution and the interference of lipid signals (**Fig. S9**). With the homogenization of sample refractive index after lipid removal, deep VISTA imaging throughout a 1 mm hippocampal tissue (effectively 250 μm in unexpanded tissues) is also achieved (**Fig. 2m**). We note that our current imaging depth is mainly limited by the short working distance of the signal-collecting condenser and could be significantly improved by replacing both the objective and the condenser with long working distance objectives (e.g., 8 mm), specifically designed for tissue-clearing imaging of whole mouse brain hemispheres^26,27^.

Since VISTA distinctly delineates the shapes of neuronal cells, processes, and likely blood vessels (**Fig. 2h-l**), we set out to validate the identities of these protein-rich biological components in VISTA images with established immunofluorescence. We were able to nicely correlate almost all structures shown in VISTA with fluorescence targets across various regions in brain tissues, including hippocampus and cortex. First, each vessel-like structure in VISTA is confirmed by lectin-DyLight594 staining, including those in vessel heavy regions within the hippocampus **(Fig. 3a-b, S10a-c)** and larger ones likely of arteries^20,28^ **(Fig. 3c-d)**. In addition to capturing vessel structures and distributions, VISTA could also image protein abundant red blood cells inside vessels that are retained in the polymer gel network (**Fig. 3c, Fig S10d-f**). Second, all the nuclei shown in VISTA have one-to-one correspondence to DAPI labeling **(Fig. 3e&g)**. We found that a small portion of these nuclei are from vascular endothelial cells, confirmed by their co-localizations with lectin vessel staining (**Fig. 3e-h,** arrowed) and GLUT1 immunostaining (**Fig. S11**, arrowed). The rest of the nuclei should come from various types of cells including neurons, astrocytes, oligodendrocytes, etc. Third, together with nuclear structures, some cells also present clear contrast of cytoplasm. Our further correlative imaging identified that all these cell bodies captured by VISTA belong to matured neuronal cells, but not to astrocytes or oligodendrocytes. This is confirmed by the co-localizations of VISTA cell bodies with immuno-fluorescence-stained NeuN (matured neuron marker, **Fig. 3i-j**) and MAP2 (marker of neuronal perikarya and dendrites, **Fig. 3k-l**); and the lack of cross-localizations to glial fibrillary acidic protein (GFAP, astrocyte cellular maker, **Fig. S12a-f**) and myelin basic proteins (MBP, oligodendrocyte cellular marker, **Fig. S12g-l**). These results also suggest that maturated neuronal cells are more protein abundant in the cytosols compared to astrocytes and oligodendrocytes, a result difficult to quantify by other methodologies. Fourth, in addition to cell bodies, neuronal processes visualized in VISTA could be assigned to dendrites, which showed decent overlap with those imaged by MAP2 stains (**Fig. 3k-l**). Our characterizations by correlative immunofluorescence imaging hence confirm that VISTA offers holistic mapping of nuclei, vessels, and neuronal cell distributions and dendritic connections in brain tissues, without any labels.

**Figure 3.**
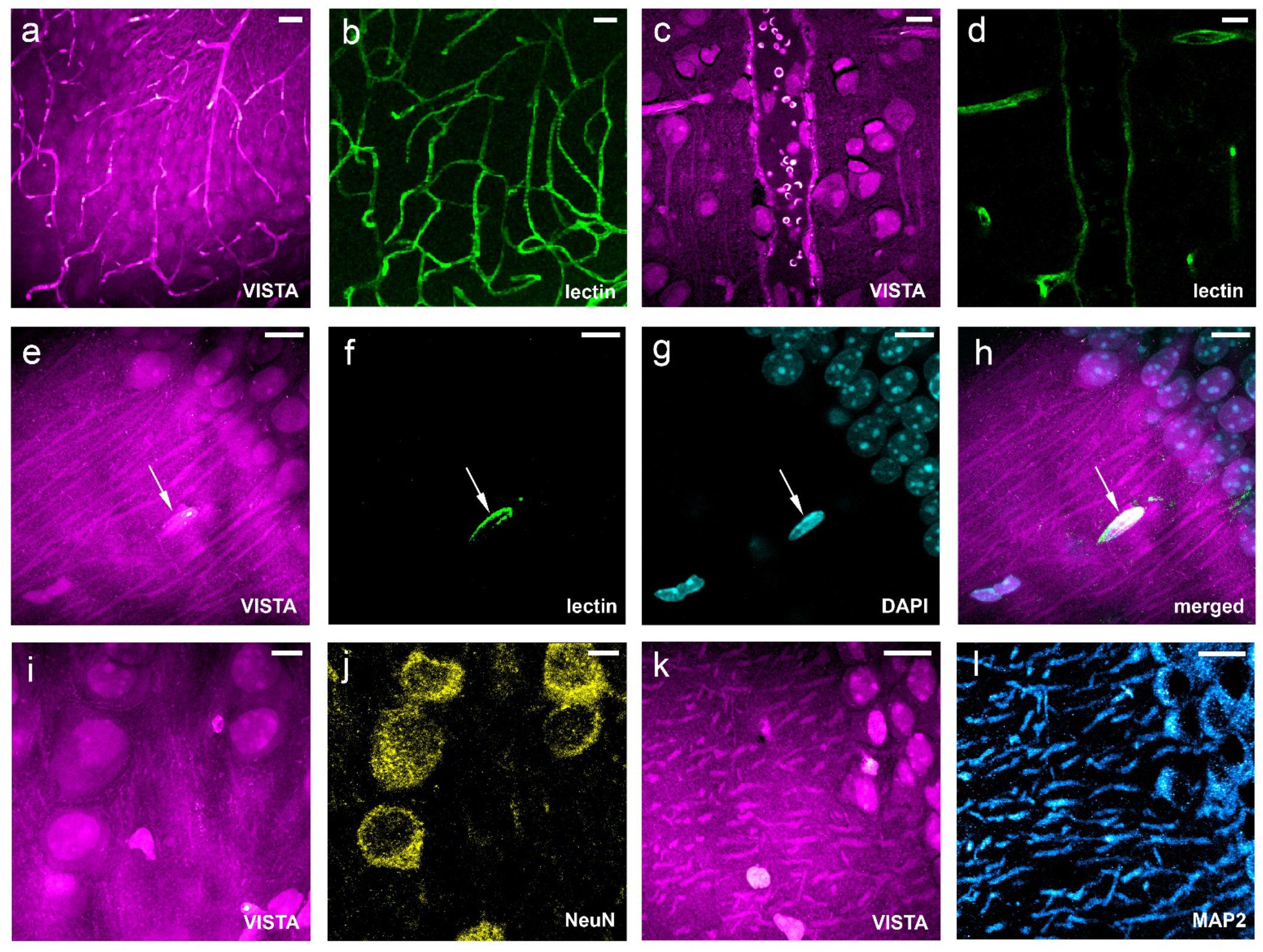
Validation of VISTA imaging features with fluorescent markers on mouse brain tissues. **a-d**, Parallel images of VISTA (a, c) and fluorescence from lectin-DyLight594 stained blood-vessel (b, d) in vessel-abundant regions. **e-g**, Parallel images of VISTA (e) with two-color fluorescence from lectin- DyLight594 stained vessels (f) and DAPI-stained nuclei (g). **h.** Three-channel merged image from e-g. **i-j**, Parallel images of VISTA (i) and fluorescence from immuno-stained NeuN, the matured neuron marker (j). **k-l**. Parallel images of VISTA (k) and fluorescence from immuno-stained MAP2, the neuronal cell body and dendrite marker (l). All images are shown as maximum intensity projection from a stack of volume images. Scale bars: 40 μm. The length scale is in terms of distance after sample expansion.

High-resolution 3D mapping of the intricate cellular, vasculature, and connectivity network in brain has been a long-sought goal for super-resolved fluorescence microscopy. Imaging such multi-cellular interplay with high throughput would significantly maximize the information value and open new avenues for versatile biological investigations, such as in stroke models and neurodegenerative diseases^25,29–31^. As we were able to assign the origins of protein-abundant structures in VISTA to specific protein targets, we then aimed to transform each identified biological structure in **Fig. 3** into individual component for multitarget analysis through image segmentation. Recently, CNN based deep learning has been implemented for SRS imaging, but mostly focused on denoising and diagnosis prediction^32–35^. The imaging segmentation requires hyperspectral SRS and yet offers limited resolution^33^. Here, we adapted a U-Net based architecture^17,36^ and trained our model with parallel VISTA and fluorescence images as input datasets for high-resolution image segmentation. VISTA images (**Fig. 4a, d, g**) were then passed through the trained model, and successfully generated predicted structures (**Fig. 4c, f, i**) that correlate well with fluorescence-labeled ground truth (**Fig. 4b, e, h**) for blood vessels (**Fig. 4c, v-lectin**), nuclei (**Fig. 4f, v-DAPI**), MAP2- immunolabeled neuronal cell bodies and dendrites (**Fig. 4i, v-MAP2**), and NeuN-immunolabeled mature neurons (**Fig. S13**). The quality and contrast of these predicted images are close to the corresponding fluorescence images. The prediction performance is quantified by the Pearson correlation coefficient (**Fig. S14**). We note that the relatively low correlation values for NeuN prediction were mainly due to low fluorescence signals obtained, likely caused by the loss of NeuN epitope during protein denaturation. With these 4 individual components successfully predicted, 4-color multiplex imaging is readily obtainable in 3D (**Fig. 4j and Fig. S15**). For more integrated insights of biological organizations, 6-to-7-component imaging could also be achieved on the same sample with additional 2 to 3 fluorescent colors (**Fig. 4k and Fig. S16**). Comparing to conventional label-free imaging, deep learning equipped VISTA offers desired target specificity for multiplex structural analysis. Comparing to sample-expansion fluorescence microscopy, which typically requires week-to-month long sample preparation with immunofluorescence^4,6,20,21^, sample-processing steps for VISTA are complete within 48 hours (**Fig. S17**). Such high-throughput nature of label-free VISTA imaging by omitting the multi-round immunostaining processes for multi-component investigations would largely facilitate our understanding of the intricate relationship between these cellular and sub-cellular structures deep inside tissues.

**Figure 4.**
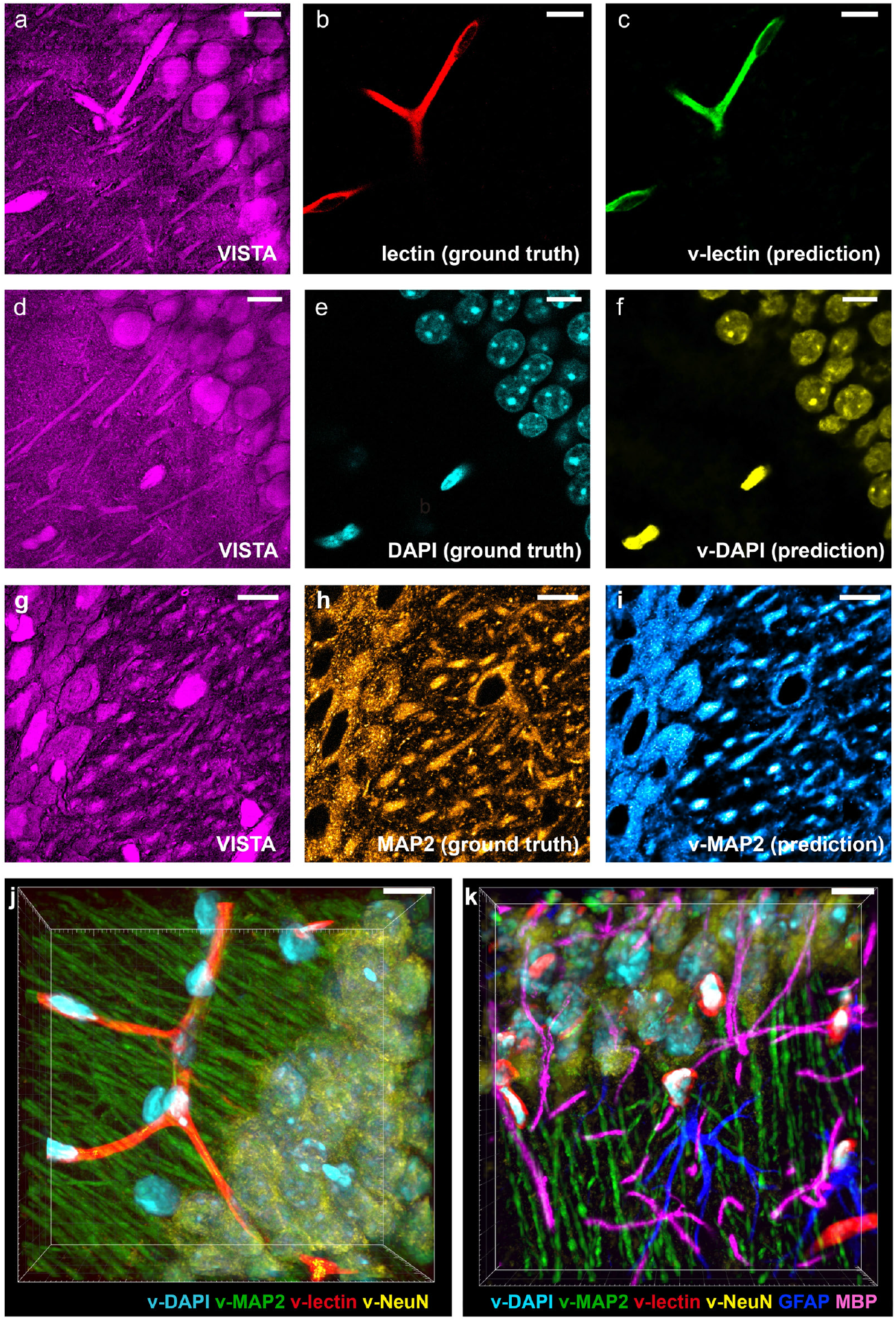
Label-free VISTA prediction for specific and multi-component imaging of brain hippocampal tissues. **a-c**, The input VISTA image (a), the ground truth fluorescence image of lectin- DyLight594 stained blood vessels (b) and the predicted VISTA-lectin (v-lectin) image of blood vessels (c). **d-f**, The input VISTA image (d), the ground truth fluorescence image of DAPI stained nuclei (e) and the predicted VISTA-DAPI (v-DAPI) image of nuclei (f). **g-i**, The input VISTA image (g), the ground truth immunofluorescence image of MAP2 stained neuronal cell bodies and dendrites (h) and the predicted VISTA-MAP2 (v-MAP2) image of neuronal cells and dendrites (i). **j**, 4-color volume imaging from label- free VISTA prediction for vessels (v-lectin, red), nuclei (v-DAPI, cyan), neuronal cell bodies and dendrites (v-MAP2, green) and matured neuron cell bodies (v-NeuN, yellow). **k**, Tandem 6-color volume imaging from label-free 4-color VISTA prediction and parallel two-color immuno-fluorescence images of GFAP (blue) and MBP (magenta). Scale bars: 40 μm. The length scale is in terms of distance after sample expansion.

## Discussion

In summary, we established VISTA as a robust and general label-free method for resolving proteinrich cellular and subcellular structures in 3D cells and tissues with an effective imaging resolution down to 82 nm. Targeting the CH_3_ vibrational groups from endogenous proteins, VISTA is free from probe bleaching, decay or quenching caused by photo-illumination or gel polymerization and hence suited for repetitive and quantitative interrogations. Implemented with machine learning, VISTA allows specific and multi-component imaging of nuclei, blood vessels, matured neuronal cells and dendrites in brain tissues. VISTA avoids low-efficiency, inhomogeneous delivery, and high cost of fluorescent antibodies, and thus offers fast throughput, uniform imaging throughout tissues, and cost-effective sample preparation, which scales up better with large human brain samples for future clinical investigations.

A few further technical improvements could be explored to bring VISTA a step forward. It could be coupled with all existing instrument-based high-resolution vibrational imaging techniques for further improvement in resolution. In particular, with a recently reported SRS configuration of frequency-doubled (i.e. wavelength-halved) excitation lasers^37^, VISTA should further push the obtainable resolution down to 30 nm. As SRS signals scale nonlinearly with the laser frequency, frequency doubling would allow a 16-time improved sensitivity to resolve lower-protein-abundance structures. In addition to imaging proteins, VISTA could be extended to imaging other types of biomolecules including DNA, RNA, lipids with proper development of molecular anchoring chemistry during sample embedding^38^. With all these features, VISTA should find a wide range of applications for mapping subcellular architectures, cell distributions and connectivity across various molecular and resolution scales in complex tissues.

## Acknowledgements

We thank Xun Wang and Dr. Lilien Voong for fruitful discussions. We are grateful to Can Li and Prof. Marianne Bronner for sharing the zebrafish embryo slices. Chenxi Qian acknowledges the support of the Natural Sciences and Engineering Research Council of Canada (NSERC Postdoctoral Fellowship). Lu Wei acknowledges the support of National Institutes of Health (NIH Director’s New Innovator Award, DP2 GM140919-01), Amgen (Amgen Early Innovation Award) and the start-up funds from California Institute of Technology.

## Data availability

The authors declare that all data supporting the findings of the present study are available in the article and its supplementary figures and tables, or from the corresponding author upon request.

## Code availability

MATLAB code used for PSF determination and Python code for U-Net training and prediction in this paper is available at https://github.com/Li-En-Good/VISTA.

