## Supplementary Information for "Super-Resolution Label-free Volumetric Vibrational Imaging"

### General Methods

**Stimulated Raman scattering (SRS) Microscopy.** A picoEmerald laser system (Applied Physics & Electronics) is used as the light source for SRS microscopy. It produces 2 ps pump (tunable from 770 nm – 990 nm, bandwidth 0.5 nm, spectral bandwidth ~ 7 $cm^{-1}$) and Stokes (1031.2 nm, spectral bandwidth 10 $cm^{-1}$) beams with 80 MHz repetition rate. Stokes beam is modulated at 20 MHz by an internal electro-optic modulator. The spatially and temporally overlapped pump and Stokes beams are introduced into an inverted multiphoton laser scanning microscope (FV3000, Olympus), and then focused onto the sample by a 25X water objective (XLPLN25XWMP, 1.05 N.A., Olympus) for imaging. Transmitted Pump and Stokes beams are collected by a high N.A. condenser lens (oil immersion, 1.4 N.A., Olympus) and pass through a bandpass filter (893/209 BrightLine, 25 mm, Semrock) to filter out Stokes beam. A large area (10×10 mm) Si photodiode (S3590-09, Hamamatsu) is used to measure the pump beam intensity. A 64 V reverse-bias DC voltage is applied on the photodiode to increase saturation threshold and reduce response time. The output current is terminated by a 50-Ω terminator and pre-filtered by a 19.2-23.6-MHz band-pass filter (BBP-21.4+, Mini-Circuits) to reduce laser and scanning noise.

The signal is then demodulated by a lock-in amplifier (SR844, Stanford Research Systems) at the modulation frequency. The in-phase X output is fed back to the Olympus IO interface box (FV30-ANALOG) of the microscope. Image acquisition speed is limited by 30 μs time constant set for the lock-in amplifier. Correspondingly, we use 80 μs pixel dwell time, which gives a speed of 21s per frame for a 512-by-512-pixel field of view. Pump laser is tuned to 791.3 nm for imaging protein $CH_3$ vibrational mode at 2940 $cm^{-1}$. Laser powers on sample are measured to be 30 mW for Pump beam and 200 mW for modulated Stokes beam. 16-bit grey scale images are acquired by Olympus Fluoview 3000 software. Volumetric images were acquired by collecting a z-stack with a step size of 1 micron in z direction.

**Reagents and materials.** Sodium acrylate (SA, Sigma-Aldrich), acryl amide (AA, Sigma-Aldrich), N,N′-methylenebisacrylamide (BIS, 2%; Sigma-Aldrich), ammonium persulfate (APS, Sigma-Aldrich), tetramethylethylenediamine (TEMED, Sigma-Aldrich), sodium dodecyl sulfate (SDS), Triton X-100, Tween-20, and deuterium oxide were obtained from Sigma-Aldrich, and 1.0 M Tris was obtained from Biosolve. Nuclease-free water was purchased from Ambion–Thermo Fisher. Acrylamide (2,3,3-D3) was obtained from Cambridge Isotope Laboratories. Deuterated sodium acrylate was prepared from acrylic acid (2,3,3-D3, Cambridge Isotope Laboratories) and sodium hydroxide (Sigma-Aldrich). DAPI was purchased from Thermo Fisher (D1306, Thermo Fisher). Primary antibodies: anti-myelin basic protein in rat (Abcam, ab7349); anti-myelin basic protein in rabbit (Abcam, ab40390); anti-GFAP in chicken (Abcam, ab4674); anti-chicken IgY, Alexa 488 (Invitrogen, A-32931); anti-GFP Alexa Fluor 647 (Invitrogen, A-31852); anti-MAP2 (Cell Signaling Technology,8707); anti-NeuN (Cell Signaling Technology, 24307); anti-GLUT-1 (Millipore Sigma, 07-1401); Lycopersicon Esculentum Lectin DyLight®594 (Vector Laboratories, DL-1177-1). Secondary antibodies: goat anti-rat IgG, Alexa Fluor 568 (Invitrogen, A-11077); goat anti-mouse IgG, Alexa Fluor 647 (Invitrogen, A-21236); goat anti-rabbit IgG, Alexa Fluor 488 (Invitrogen, A-11034); goat anti-chicken IgY, Alexa Fluor 647 (Invitrogen, A-21449).

**Hydrogel embedding: gelation, denaturation and expansion.** Stock solutions include an incubation solution (30% AA in 4% PFA), and a gelation solution (7% SA, 20% AA, 0.1% BIS in 1x PBS) were made and stored in 4 °C and -20 °C, respectively. The free-radical initiator APS and accelerator TEMED were dissolved and diluted in nuclease-free water to a concentration of 10% (w/w), and stored at -20 °C as stocks. Prior to a typical hydrogel embedding step, the cell or tissue samples were incubated in a solution of 30% AA in 4% PFA under different conditions depending on the sample type (detailed in the following sections). The gelation solution, the free-radical initiator and accelerator were thawed and kept at 4 °C before the gelation step. Coverslips with the cell or tissue samples were placed at the bottom of a pre-fabricated and pre-cooled gelation chamber, with the sample facing upward. After adding sufficient amount of gelation solution (7% SA, 20% AA, 0.1% BIS in 1x PBS) to fully immerse the sample, a layer of flat Parafilm covered coverslip was placed on top of the chamber as the lid. The chamber was kept at 4 °C for 1 min as the free-radical polymerization proceeded, before it was transferred to a humid incubator for a following incubation at 37 °C for 1 h. Coverslips with gels were then incubated in sufficient amount of denaturing buffer (200 mM SDS, 200 mM NaCl, and 50 mM Tris in nuclease-free water, pH 8) in petri dishes for 15 min at room temperature. The volume of the denaturing buffer and the size of the petri dishes used were determined by the sample size and thickness after denaturation. In 15 min, gels would be detached from the coverslips. They were transferred into 1.5-ml Eppendorf centrifuge tubes filled with denaturing buffer and were incubated under different conditions depending on the sample type (detailed in the following sections). Initial expansions were carried out immediately after the denaturation step at ambient temperature, in $H_2O$ or $D_2O$, which was changed twice in 1 h. The expanded gel was then kept in $H_2O$ or $D_2O$ overnight and stores in dark. The gel expanded 4.0±0.22 times in our experiments.

**Cultured HeLa cell experiments.** In mammalian cell experiments, cultured HeLa-CCL2 (ATCC) were seeded onto coverslips (12mm, #1.5, Fisher) for 24 h. Cells were first grown in regular DMEM medium supplemented with 10% FBS and 1% penicillin-streptomycin antibiotics until they reached 70-90% confluency. Coverslips with HeLa cells were incubated in a solution of 4% PFA with 30% AA in PBS for 7–8 h at 37 °C, without normal fixation with PFA. Hydrogel embedding, including gelation, denaturation and expansion, was processed in above-mentioned steps. The denaturation after transfer into Eppendorf centrifuge tubes was at 95 °C for 30 min.

**Normal brain tissue experiments**. Mouse brains tissues harvested from mice were washed once with DPBS on ice and immediately incubated in a solution of 4% PFA with 30% AA in PBS for 24-30 h at 4 °C, then transferred to a shaker and further incubated for 12 h at 37 °C with gentle shaking. After the incubation step the tissue sample was cut into 100 μm to 250 μm thin slices (and thick slices up to 600-650 μm, for thick brain tissue experiments) using a Leica VT 1200S vibratome. Hydrogel embedding, including gelation, denaturation and expansion, was processed in above-mentioned steps. The denaturation after transfer into Eppendorf centrifuge tubes was first at 70 °C for 3 h and then at 95 °C for 1 h.

**Zebra fish embryo experiments.** Fresh zebra fish embryo samples were embedded in gelatin and snap frozen in liquid nitrogen and stored at -80 °C until ready for sectioning. Immediately before sectioning, allow the whole gelatin embedded sample warm up to -30 °C for 10 min in the cryostat. Cryosectioning was carried out on the whole frozen block, and sectioned slices were collected onto the glass slides at room temperature. The glass slides with collected sample slices were stored at -20 °C until ready for de-gelatinization and hydrogel embedding. When ready, the slides were warmed up to room temperature, and then de-gelatinized by incubation in PBS at 42 °C for 30 min. The de-gelatinized samples were washed in PBS with 0.1% Tween-20 before processing. Hydrogel embedding, including gelation, denaturation and expansion, was processed in above-mentioned steps. For thin (50-μm) slices of the zebra fish embryo we used, the denaturation after transfer into Eppendorf centrifuge tubes was at 95 °C for 30 min.

**Thick brain tissue experiments**. Thick brain tissues were cut to slices with thickness of at least 250 μm (1000 μm after expansion). PFA+AA incubation and hydrogel embedding, including gelation, denaturation, and expansion, was processed in above-mentioned steps similarly to the normal tissue processing. The denaturation after transfer into Eppendorf centrifuge tubes was first at 70 °C for 3 h and then at 95 °C for 4h. Thicker tissue can also be imaged by VISTA with longer 95 °C denaturation time.

**Immunostaining**. In an immuno-labeling process, 60-μm- to 150-μm-thick mouse brain coronal slices were embedded in a hydrogel, denatured, pre-incubated with PBS with 1% (wt/vol) Triton X-100 (PBST) for 15 min, and subsequently incubated with primary antibodies at a typical 1:100 dilution with PBST at 37 °C for 16h, followed by washing with PBST three times at 37 °C for 1-2 h. The samples were then incubated with secondary antibodies at a 1:100 dilution with PBST at 37 °C for 12-16 h, followed by washing with PBST three times at 37 °C for 1-2 h.

**Sample mounting and imaging**. Expanded cell or tissue samples were kept in $D_2O$ for imaging. Grace Bio-Labs Press-To-Seal silicone isolators with appropriate opening sizes and depths were used as spacers between microscope slides (1 mm, VWR) and coverslips (12 mm, #1.5, Fisher). In particular, the thick brain tissue samples (after expansion) were placed in a 4-mm silicon isolator to avoid any pressure and damage to the sample. For control experiments on normal PFA fixed HeLa cells, 0.5-mm Press-To-Seal silicone isolators or the common Grace Bio-Labs SecureSeal™ spacers were used. Confocal images were obtained by the Olympus FluoView™ FV3000 confocal microscope with SRS setup described above.

**SRS imaging of beads.** Polystyrene beads (0.1 μm mean particle size, Sigma-Aldrich, Inc.) were resuspended in deionized water by a 1:2000 dilution. The resuspension step required vortexing and sonication for 20 min at room temperature. Before SRS imaging, the diluted beads suspension was sealed between a glass slide and a coverslip, which was then stored in dark and left to settle overnight.

**Image processing and data analysis**. Images color-coding and intensity profile were done by ImageJ. Intensity normalization of the z-stacks was done in ImageJ. 3D rendering of z-stacks was done in Imaris View. Data plotting and analysis, including spectral plots and Gaussian fitting were performed in OriginLab.

**Calculating the resolution of SRS microscopy and the VISTA method.** To determine the point spread function (PSF) of the imaging system, deconvolution of the actual size of the beads was done by simulations in MATLAB 2019b. Line profile of the bead image was fitted by Gaussian approximation and bead object was modelled as a circle.

**Fluorescence imaging.** The fluorescence images of processed samples with fluorescent labels were obtained with a 25×, 1.05 NA water-immersion objective with the Olympus Fluoview system. Single-photon confocal laser scanning imaging was performed with 405-, 488-, 561-, and 640-nm lasers (Coherent OBIS). The images were visualized and analyzed with Fiji or Imaris Viewer.

**U-Net construction, training and prediction for label-free imaging.** The prediction of subcellular structures from SRS images was based on a U-Net CNN convolutional neural network demonstrated by Ounkomol et al [1]. Training data were collected by sequentially acquiring respective fluorescence targets and protein SRS images on the same field of view. Image sets were generated by performing a z scan with a z-direction step size of 1 micron. Such a step size is larger than the axial resolutions of both fluorescence and SRS. Before training, the fluorescence and SRS images were background subtracted in imageJ with a rolling ball radius of 50 pixels (0.497 micron/pixel) before training. The image sets were separated at a 1:3 ratio for testing and training set, respectively. The models were trained by batches of 128 pixels x 128 pixels patches subsampled from the original images. The training was performed using the Adam optimizer to optimize the mean squared error between the fluorescence image and the predicted image. The learning rate set at 0.001 and trained for 50,000 epochs with batch size of 32 images. All the trainings and predictions were run on a node of High Performance Computing Center at Caltech equipped with Nvidia P100 GPU containing 16 GB of memory.

**Model accuracy.** Model accuracy was quantified by the Pearson correlation coefficient:

$$r = \frac{\sum(x-\overline{x})\,(y-\overline{y})}{\sqrt{\sum(x-\overline{x})^2\sum(y-\overline{y})^2}}$$

between the pixel intensities of the model's output, y, and independent ground-truth test images, x, for all the images of the test sets except for lectin. The Pearson correlation coefficient for lectin was calculated for images with lectin signal, since the random noise in the background could not be predicted by the model.

**Data and code availability.** The authors declare that all data supporting the findings of the present study are available in the article and its supplementary figures and tables, or from the corresponding author upon request. MATLAB code used for PSF determination and Python code for U-Net training and prediction in this paper is available at https://github.com/Li-En-Good/VISTA.

**Animals.** All animal procedures performed in this study were approved by the California Institute of Technology Institutional Animal Care and Use Committee (IACUC), and we have complied with all relevant ethical regulations. The C57BL/6J (000664), Thy1-YFP (003709) mouse lines used in this study were purchased from the Jackson Laboratory (JAX) and bred in our animal facility. 6- to 8-week-old, male and female C57BL/6J, Thy1-YFP mice were used for tissue collection. At the day of collection, the mice were anesthetized with Euthasol (pentobarbital sodium and phenytoin sodium solution, Virbac AH) and transcardially perfused with 30–50 mL of 0.1 M phosphate buffered saline (PBS) (pH 7.4). After this procedure, the brains were harvested and proceeded to VISTA processing.

[1] C. Ounkomol, S. Seshamani, M. M. Maleckar, F. Collman, and G. R. Johnson, "Label-free prediction of three-dimensional fluorescence images from transmitted-light microscopy," *Nat. Methods* 15(11), 917–920 (2018).

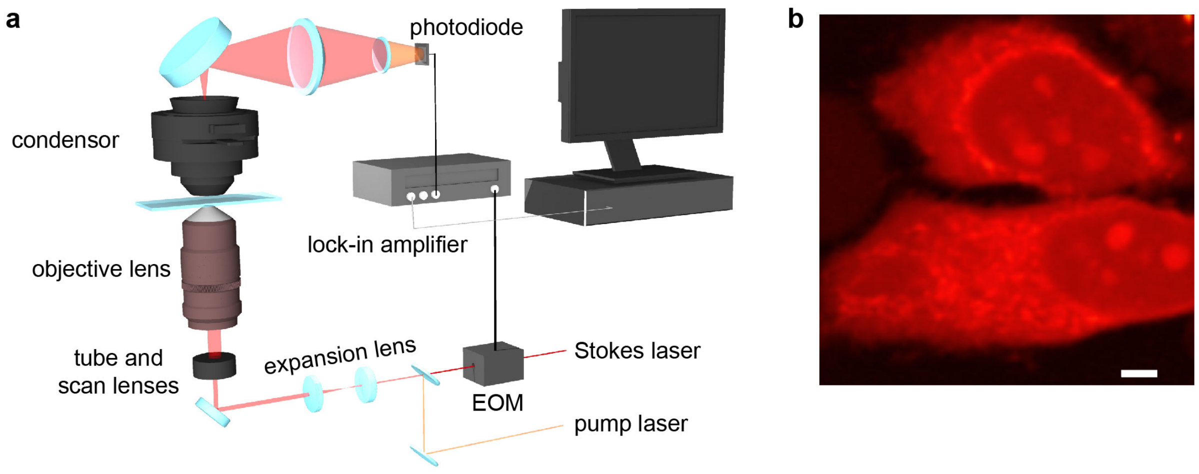


**Figure S1. Setup for the stimulated Raman scattering (SRS) microscopy.** (a) Instrumental setup of SRS microscope. EOM: Electro-Optic Modulator. When the energy difference between the Pump laser photons and the Stokes laser photons matches with the vibrational frequency of target chemical bonds (e.g. 2940 $cm^{-1}$ for the symmetric vibrational motion of $CH_3$), the chemical bonds are efficiently driven from the vibrational ground state to the vibrational excited state, creating stimulated Raman loss in pump beam, which is subsequently detected by a photodiode and provides the imaging contrast. (b) A representative regular-resolution SRS image of HeLa cells targeting the protein $CH_3$ vibration at 2940 $cm^{-1}$. Scale bar: 3 μm.

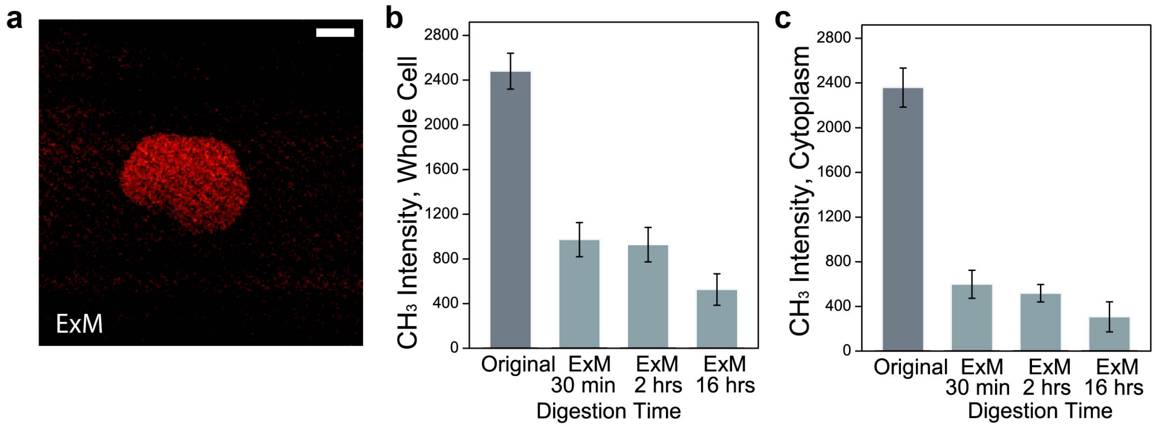


**Figure S2. SRS quantification of protein loss following the ExM protocol.** (a) Contrast-enhanced SRS image for Fig. 1c with a 2-time enhancement on the intensity scale. Scale bar: 20 μm. The length scale is in terms of distance after sample expansion. (b)-(c) Quantification of protein retention level on different protease digestion time by proteinase K on whole cells (b) and from the cytoplasm (c). SRS images of protein $CH_3$ from HeLa cells were acquired after designated digestion times. The protein signals decrease rapidly upon protease treatment, confirming that the digestion step in ExM causes significant protein loss. Error bar: SD.

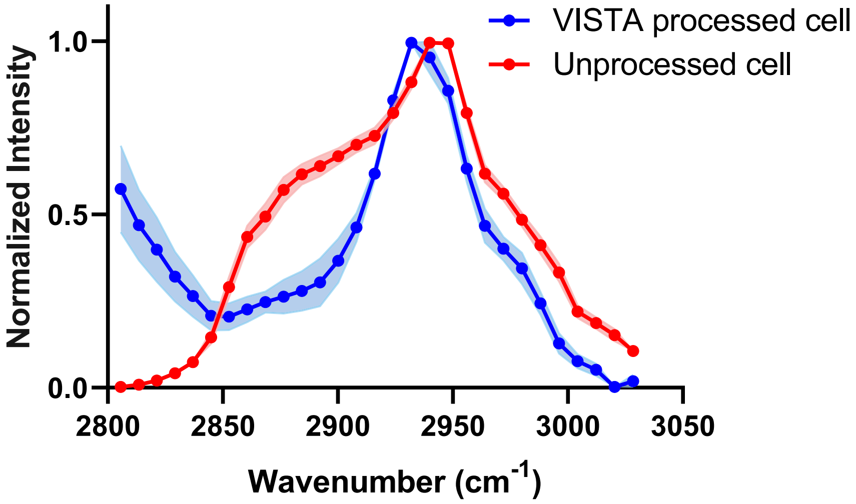


**Figure S3. Hyperspectral SRS from HeLa cells with and without VISTA processing.** The spectra were acquired by sweeping pump laser wavelength with 0.5 nm step size from 783.3 to 799.8 nm, with Stokes laser fixed at 1031.2 nm. The evident loss of lipid peak at 2855 cm$^{-1}$ between normalized SRS spectra from unprocessed cells (red) and VISTA processed cells (blue) confirms the loss of lipid content from the protein denaturation step, which washes out the lipids. The intact peak shape for $CH_3$ at 2940 cm$^{-1}$ confirms that VISTA retains proteins. The shoulder around 2800 cm$^{-1}$ for VISTA processed cell spectrum (blue) originates from $D_2O$. VISTA processed cell, N=10; unprocessed cell, N=15. Error bar: SD

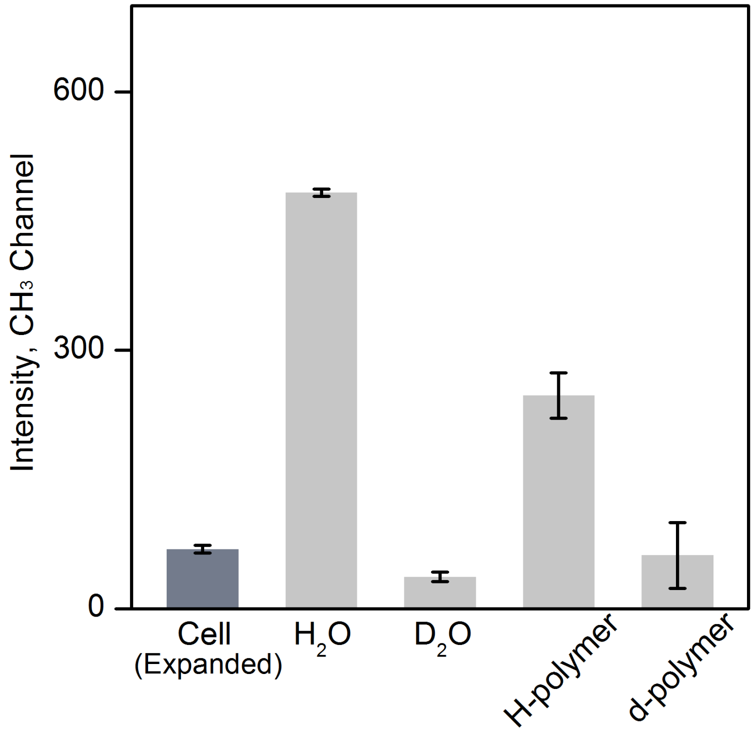


**Figure S4. Quantification of backgrounds in VISTA $CH_3$ images on HeLa cells.** Background contributions from each component (i.e. O-H stretch tail from $H_2O$ and C-H vibrations from the polymers composed of sodium acrylate and acrylamide monomers) were obtained by comparing normal VISTA cell images with VISTA images using corresponding deuterated components. The change of the backgrounds is constant across field of views after deuteration. Error bar: SD.

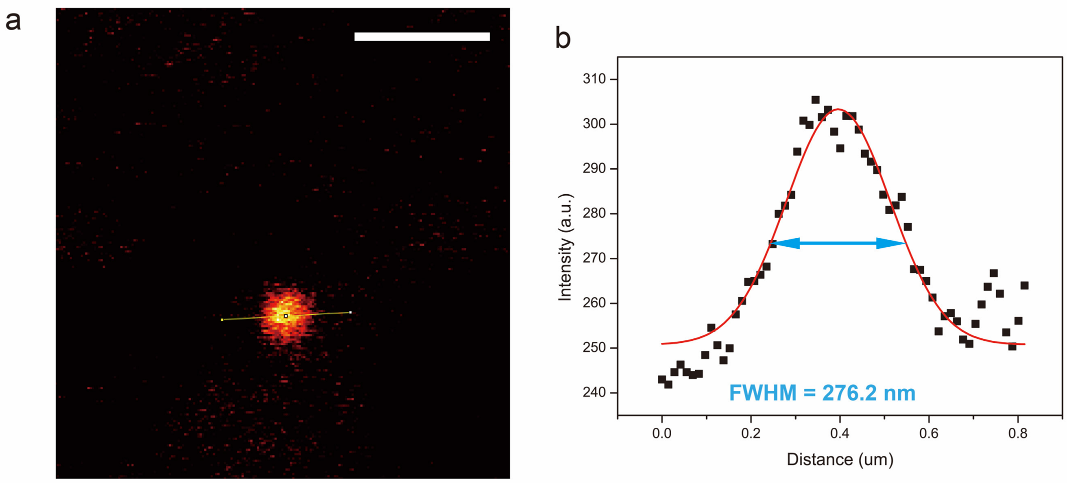


**Figure S5. Quantification of SRS resolution under a higher numerical aperture (NA) objective lens.** A representative image (a) and the corresponding fitted cross-section profile (FWHM of 276.2nm) (b) from a 100-nm polystyrene bead, by targeting the C-H vibration at 3050 $cm^{-1}$ with a 60X water objective (Olympus, UPLSAPO60XW, 1.2 NA). Scale bar: 2 μm.

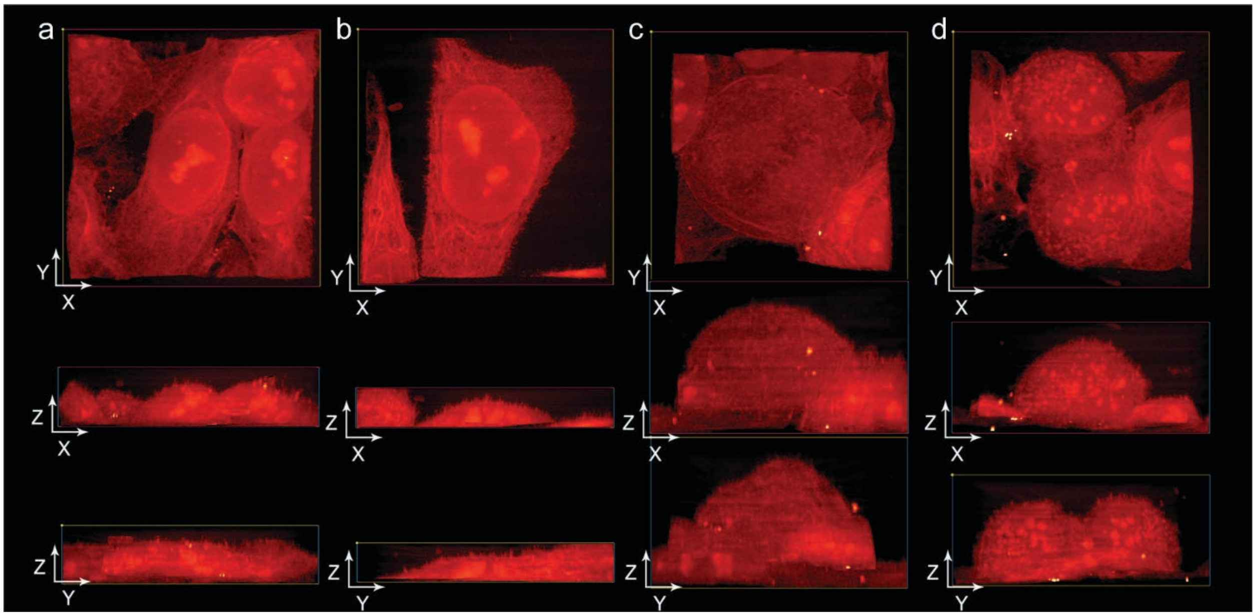


**Figure S6. High-resolution three-dimensional VISTA views of cellular morphology and subcellular structures of interphase and mitotic HeLa cells.** (a-b) Volume HeLa cells at interphase. The network structure in the cytosol is clear. In the x-z view, minor upward extrusions from cell surface (zx view) were also captured. (c-d) Volume HeLa cells during mitosis for single-z images shown in Fig. 2b and Fig. 2d.

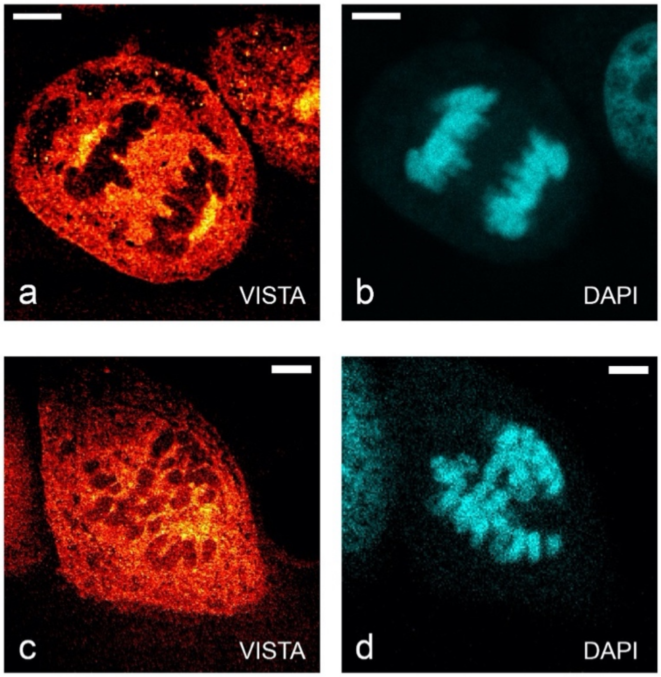


**Figure S7. Correlative VISTA images and the fluorescent images with DAPI stain on mitotic HeLa cells.** Fluorescent DAPI stains confirms that the dark regions shown in VISTA images on anaphase (a-b) and prophase (c-d) HeLa cells are chromosomes with a low $CH_3$ signals (i.e. low protein contents). Scale bars: 20 μm. The length scale is in terms of distance after sample expansion.

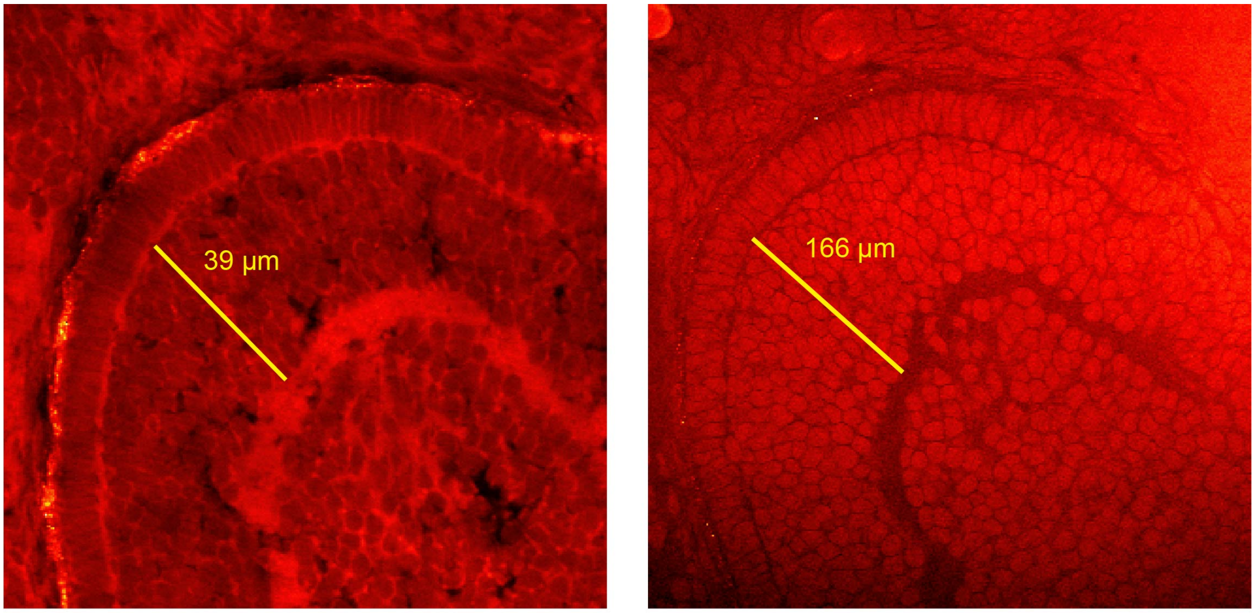


expansion ratio = 166(µm) / 39(µm) = 4.25

**Figure S8. Quantifying the expansion ratio of VISTA on tissues.** Left, before processing. Right, after expansion.

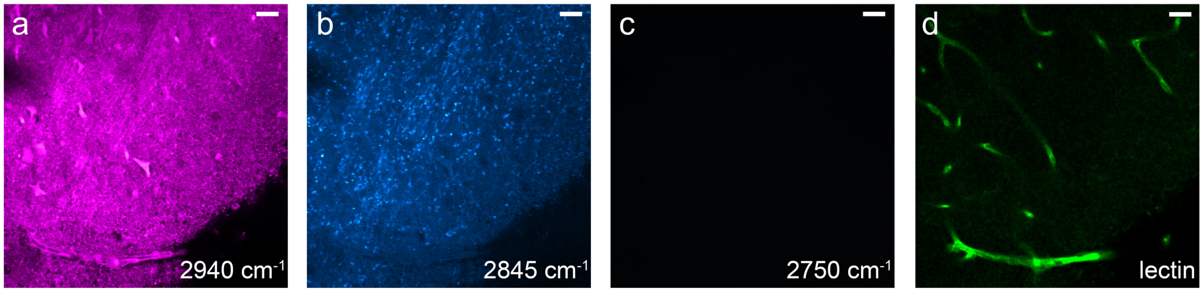


**Figure S9. Regular-resolution SRS imaging of a PFA-fixed but unprocessed mouse brain tissue.** a-c, SRS imaging at 2940 $cm^{-1}$ (a, $CH_3$), 2845 $cm^{-1}$ (b, $CH_2$), and 2750 $cm^{-1}$ (c, off-resonance image). Cells, processes and vessels could not be clearly identified from these SRS images. d, fluorescence image of lectin-stained blood vessel on the same tissue. Scale bars: 20 µm.

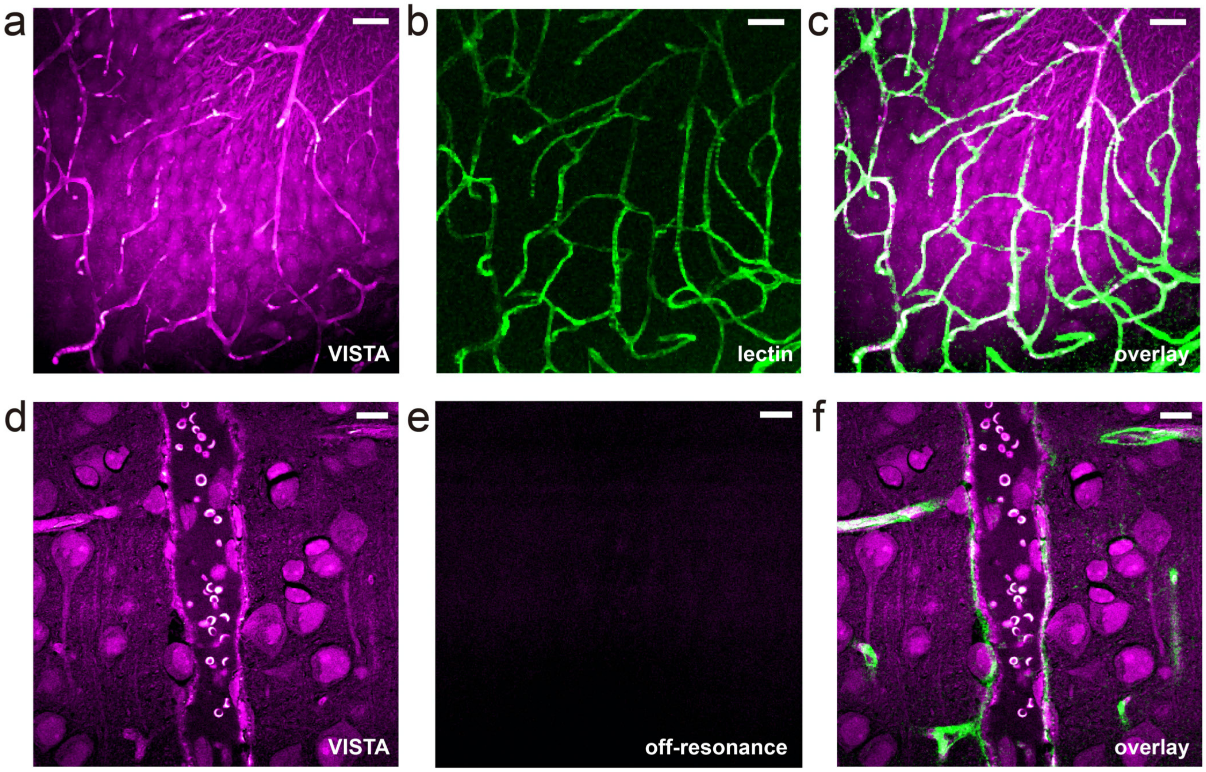


**Figure S10. Correlative VISTA images and the fluorescent images with lectin-stained vessels in mouse brain tissues.** (a-c) The overlay image (c) for parallel images of VISTA (a, shown in Fig. 3a) and fluorescence from lectin-DyLight594 stained blood-vessels (b, shown in Fig. 3b). (d-e) VISTA image of a larger vessel (likely an artery) showing red blood cells (d, shown in Fig. 3c) and the corresponding off-resonance image at 2810 $cm^{-1}$ confirms that signals in the VISTA image (including that of the red blood cells) are exclusively SRS signals. (f) The overlay image for VISTA (Fig. 3c) and lectin-DyLight594 stained fluorescence (Fig. 3d). Scale bars: 40 μm. The length scale is in terms of distance after sample expansion.

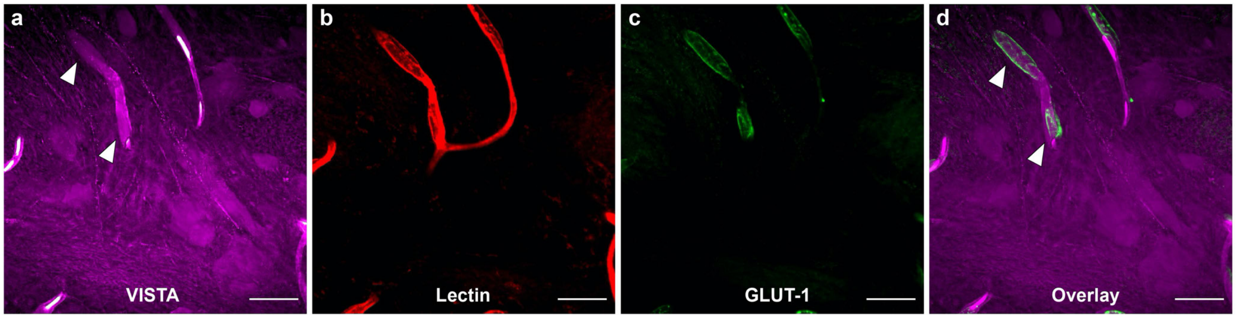


**Figure S11. Immuno-fluorescence reveals the cells in VISTA other than neuronal cells to be vascular endothelial cells.** Elongated elliptical nuclei shown in VISTA (a, white arrow-headed, VISTA) co-localize (d, Overlay) with both fluorescent lectin-stained vessels (b, Lectin), and GLUT-1 (c, glucose transporter 1, brain endothelial cell marker) immuno-fluorescence-stained brain vascular endothelial cells. Images are shown as maximum Z projection. Scale bars: 50 μm. The length scale is in terms of distance after sample expansion.

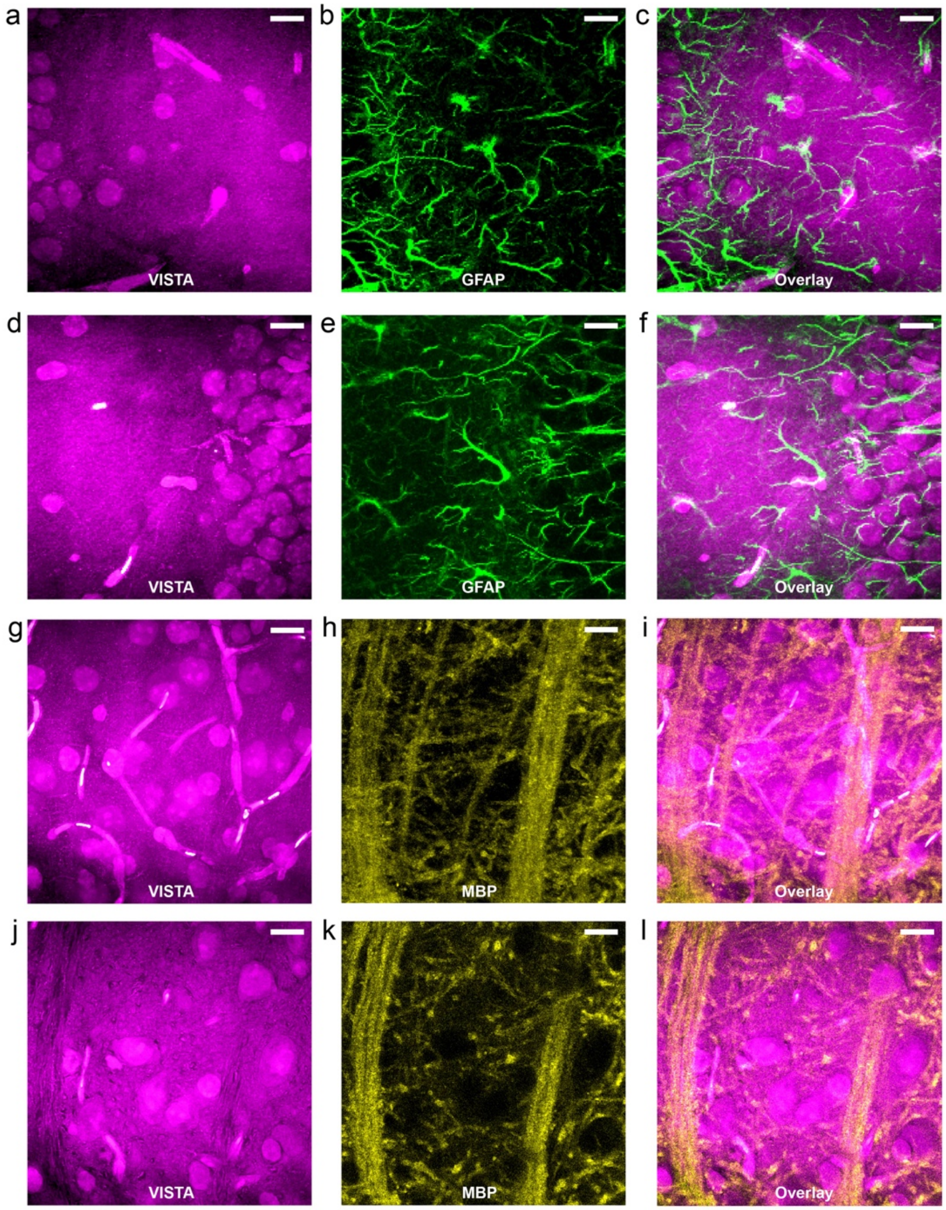


**Figure S12. Immunofluorescence confirms the absence of cytoplasmic structures from astrocytes and oligodendrocytes in VISTA.** (a-f) Maximum z projection of VISTA image (a, d), immuno-stained fluorescence image with GFAP (Glial Fibrillary Acidic Protein, the astrocyte cellular maker) antibodies (b, e) and the overlay (c, f). (g-i) maximum z projection of VISTA image (g), immuno-stained fluorescence image with MBP (Myelin Basic Protein, the myelin and oligodendrocyte cellular maker) antibodies (h) and the overlay (i). (j-l) single-slice VISTA image (j), immuno-stained fluorescence image with MBP antibodies (k) and the overlay (l). No correlation is identifiable between VISTA revealed cytoplasmic structures and GFAP or MBP stained cellular fluorescence images. Scale bars: 40 µm. The length scale is in terms of distance after sample expansion.

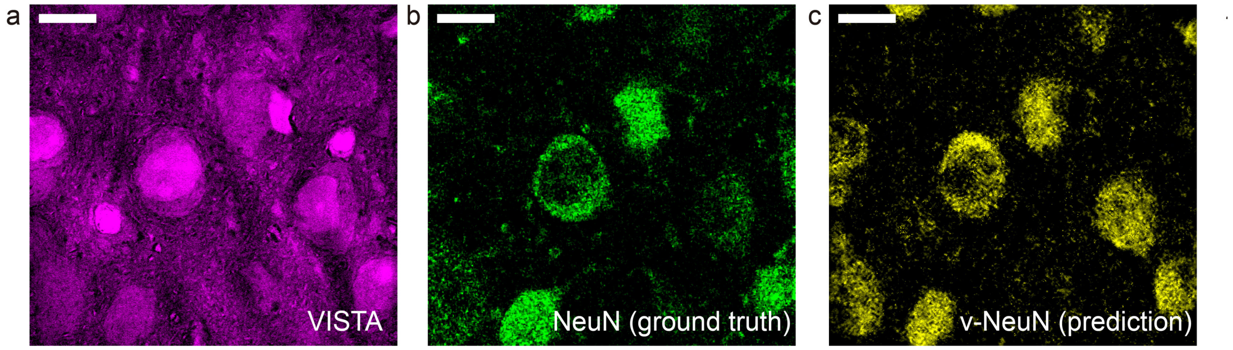


**Figure S13. Label-free VISTA prediction for neuronal cell bodies in brain tissues.** a-c, The input VISTA image (a), the ground truth fluorescence image of NeuN stained matured neurons (b) and the predicted VISTA-NeuN (v-NeuN) image of matured neurons from a (c). Scale bars: 40 μm. The length scale is in terms of distance after sample expansion.

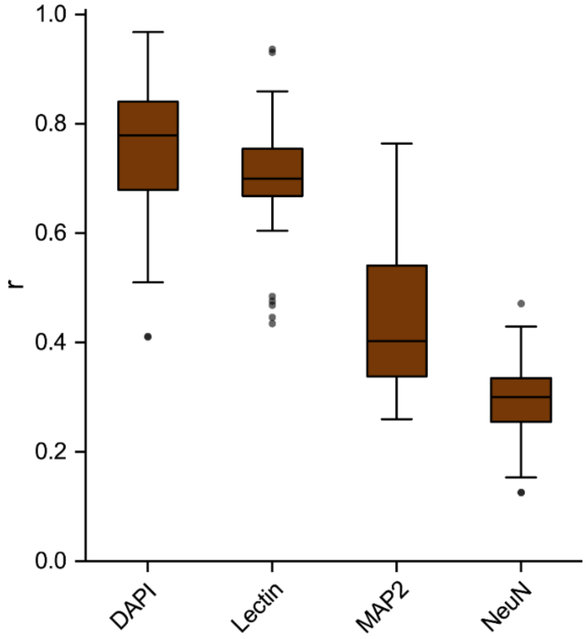


**Figure S14. Model performance for the U-Net models, quantified by Pearson correlation coefficient (r).** Each point in the plot represents a target/predicted image pair. The boxes indicate the 25th, 50th, and 75th percentile of the Pearson's r for each model, with whiskers with maximum 1.5 interquartile range. Points within the box are not shown. DAPI, N=48; Lectin, N=37; MAP2, N=54; NeuN, N=130.

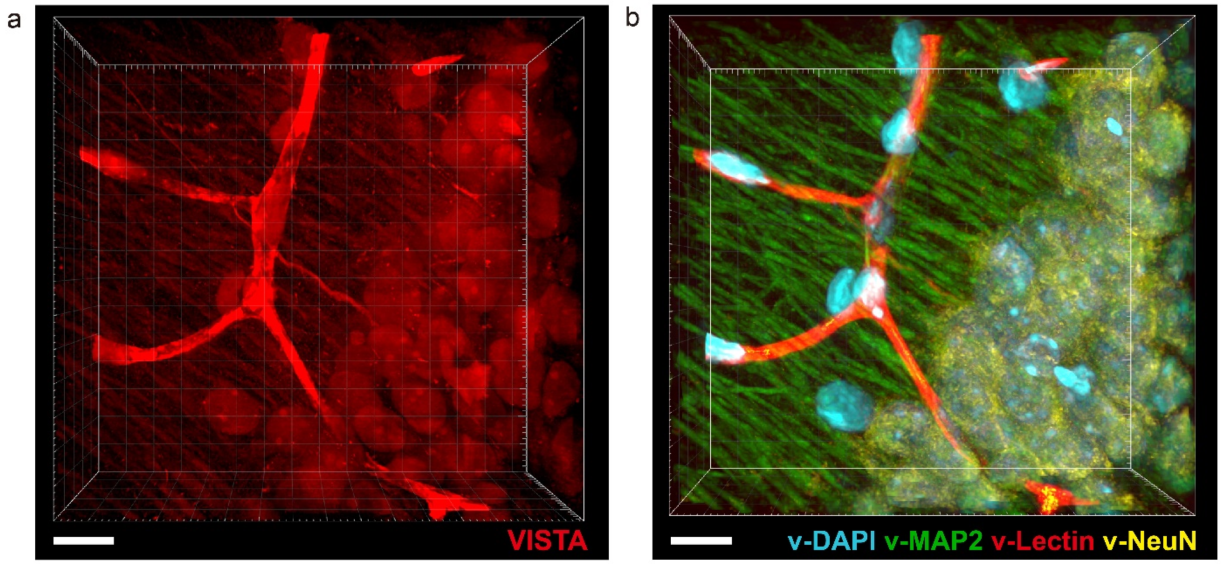


**Figure S15. Label-free VISTA prediction for multi-component imaging of mouse hippocampus.** (a) The input VISTA image; (b) the predicted multicolor image with predicted v-DAPI (cyan), v-MAP2 (green), v-lectin (red) and v-NeuN (yellow) components, as shown in Fig. 4j. Scale bars: 40 µm. The length scale is in terms of distance after sample expansion.

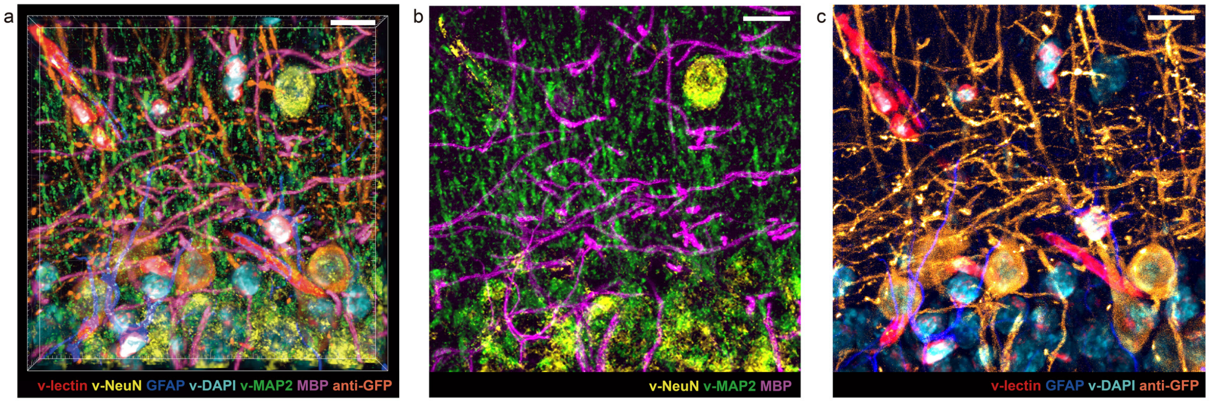


**Figure S16. 7-color multiplex imaging combining both label-free VISTA prediction and fluorescence imaging in hippocampus of Thy1-YFP mouse.** (a) Volume 3D presentation of 7-color overlay image. (b-c) Maximum z projection view of 7 components in two sets of 3-color (b) and 4-color (c) overlay. VISTA components: v-NeuN (yellow), v-MAP2 (green), v-DAPI (cyan), v-lectin (red); immuno-fluorescence components: GFAP (blue), MBP (magenta) and anti-GFP (orange). Scale bars: 40 µm. The length scale is in terms of distance after sample expansion.

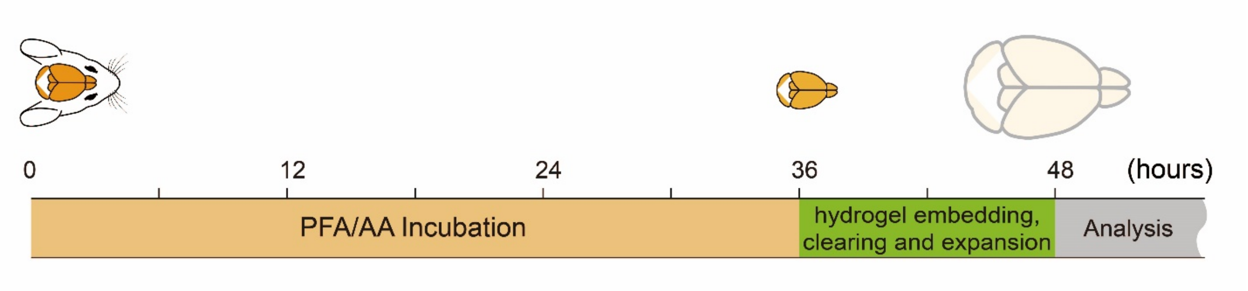


**Figure S17. Workflow of the sample processing steps for VISTA.** Typical sample processing steps were completed with a 48-hour for mouse brain tissues before VISTA analysis.
